# Distinguishing the behavioral potencies of α-pyrrolidino-phenone cathinone stimulants

**DOI:** 10.1101/2020.12.14.422779

**Authors:** Michael A. Taffe, Jacques D. Nguyen, Sophia A. Vandewater, Yanabel Grant, Tobin J. Dickerson

## Abstract

The α-pyrrolidino-phenone cathinone stimulants first came to widespread attention because of bizarre behavior consequent to the use of α-pyrrolidinopentiophenone (α-PVP, “flakka”) reported in the popular press. As with other designer drugs, diversification of cathiones has been driven by desireable subjective effects, but also by attempts to stay ahead of legal controls of specific molecules. The α-pyrrolidinohexiophenone (α-PHP) and α-pyrrolidinopropiophenone (α-PPP) compounds have been relatively under-investigated relative to α-PVP and provide a key opportunity to also investigate structure-activity relationships, i.e., how the extension of the alpha carbon chain may affect potency or efficacy. Male and female rats were used to contrast the effects of α-PHP and α-PPP with those of α-PVP in altering wheel activity and effects on spontaneous locomotion and body temperature were assessed in female rats. The α-PHP and α-PVP compounds (5, 10 mg/kg, i.p.) suppressed wheel activity in female and male rats, whereas α-PPP was only effective in female rats. Inhalation of α-PHP or α-PVP by female rats suppressed wheel activity for an abbreviated duration, compared with the injection route. Spontaneous activity was increased in a dose-dependent manner by all three compounds in female rats, and a small decrements in body temperature were observed after the highest dose of all three compounds. These data show that all three of the α-pyrrolidino-phenone cathinones exhibit significant stimulant-like activity in both male and female rats. Differences were minor and were mostly in potency and the duration of activity. Abuse liability is therefore likely to be equivalent for all three pyrrolidino-phenones.

## 1. Introduction

The synthetic cathinone drugs α-pyrrolidinopentiophenone (α-PVP) and 3,4-methylenedioxypyrovalerone (MDPV) are potent psychomotor stimulants which have been adopted by some stimulant using human populations (Adamowicz et al., 2016; Anderson, 2015; Beck et al., 2018; Benzie et al., 2011; Milian, 2015; Richman et al., 2018). These trends have been attributed initially to the lack of legal controls, but also to low cost and availability when other drugs become less readily accessed (Institoris et al., 2015), even after authorities designated them as controlled substances. MDPV was the first of these closely-related compounds to emerge in recreational use, however the α-pyrrolidino-phenone cathinone derivative stimulants came to broad public attention around 2015 (Anderson, 2015; D’Oench, 2015; Milian, 2015) because of the use of α-PVP, sometimes called “flakka” in the popular press. Early reports suggested a trend for inhaling flakka with e-cigarette devices, which introduces a further complication on assessing drug effects. The α-PVP compound was initially placed on the DEA Schedule of controlled substances in 2014 (Drug Enforcement Administration, 2014) and finalized in 2017 (Drug Enforcement Administration, 2017). Legal status has been at least a partial driver of the use and popularity of particular cathinone derivative drugs over an extended period of time. With respect to the pyrrolidino-phenones, an article in New York magazine in 2016 covering the fondness of software magnate John McAfee for α-PHP (Wise, 2016) specifically noted the fact that while related to α-PVP, it was at the time not controlled and “*so McAfee can order it with impunity from suppliers in China*”. There are also more recent suggestions from regional law enforcement that α-PHP has been used explicitly because it had been legal (Fisher, 2019). This may explain why α-PHP occurred in the top 10 phenethylamine list in the 2017, 2018 annual reports of the National Forensic Laboratory Information System (NFLIS) and in the 2019 NFLIS mid-year report, whereas α-PVP appeared up through 2018, but did not appear in the latest available report (DEA, 2019a, 2020).The α-PHP derivative was placed on the schedule temporarily in May of 2019 (DEA, 2019b) and α-PPP is currently not schedule controlled; see (Bonson et al., 2019) for review of DEA schedule of cathinone substances. Although legal status may drive initial use, these substances continue to be used after being Schedule controlled. For example there continue to be arrests all around the USA for MDPV (McGee, 2019; Swirko, 2020) or α-PVP (Fox12Staff, 2019; Keel, 2019; WALBNewsTeam, 2020) possession.

This class of pyrrolidino-phenone cathinone compounds are dopamine transporter (DAT) and norepinephrine transporter (NET) selective (over the serotonin transporter; SERT) inhibitors which do not serve as substrates/releasers (Eshleman et al., 2017) as do commonly recognized amphetamine-class stimulants such as methamphetamine and 3,4-methylenedioxy-methamphetamine and even some of the cathinones such as mephedrone or methylone (Simmler et al., 2013; Simmler et al., 2014). Cocaine is a classic non-substrate monoamine transporter inhibitor psychomotor stimulant, thus it is unsurprisingly that MDPV (Aarde et al., 2013; Aarde et al., 2015b; Gannon et al., 2017b; Watterson et al., 2014) and α-PVP (Aarde et al., 2015a; Gannon et al., 2017a; Gannon et al., 2017b; Huskinson et al., 2017; Javadi-Paydar et al., 2018a; Schindler et al., 2019) are highly effective reinforcers in animal models. MDPV and a-PVP appear to have similar potency as reinforcers (Aarde et al., 2015a), consistent with similar potencies as DAT inhibitors *in vivo* (Gannon et al., 2018). There are reasons, however, to predict that there may be differences in the behavioral and physiological effects of the three analogs in the living organism. For example, α-PHP and α-PPP differ from α-PVP in the length of the alpha carbon chain (**Figure 1**) which results in predicted lipophilicity relationships of α-PHP > α-PVP > α-PPP. Lipophilicity may affect the entry of the drug into the brain *in vivo* once ingested, producing potency differences that may not be readily predicted from an *in vitro* assay. Likewise, there is evidence that the metabolism pathway depends on the length of the alpha carbon side chain with a major difference between α-pyrrolidinobutiophenone (α-PBP) and α-PHP with α-PVP in the middle (Matsuta et al., 2018); likely α-PPP, not included in that study, would be most similar to α-PBP. Although similar potency at the DAT suggests a similar profile of acute effects of α-PHP and α-PVP, a limited study from this lab has shown that α-PHP is slightly more effective than α-PVP as a reinforcer in a self-administration (Javadi-Paydar et al., 2018b). In recent studies, rats self-administered similar numbers of infusions of α-PPP (0.1-0.32 mg/kg/infusion) and MDPV (0.05 mg/kg/infusion) in a 96 hour binge IVSA paradigm (Nagy et al., 2020), and α-PPP generated a right-shifted dose-response curve compared with MDPV or α-PVP in a 2 hr IVSA paradigm (Gannon et al., 2018). In both cases this suggests a lower potency of α-PPP relative to α-PVP, *in vivo*, as it is *in vitro* (Gannon et al., 2018; Kolanos et al., 2015). It also cannot be overlooked that off-target effects may differ across the analogs, for example Chen et all (2020) report a change in affinity for muscarinic acetylcholine receptors scales upward from α-PPP to α-PHP in association with the length of the alpha carbon chain. This association was not present in the 3,4-methylenedioxy substituted analogs, i.e., 3,4-methylenedioxypyrovalerone (MDPV) and its alpha carbon substituted analogs, cautioning further against simplistic structure-activity inferences about effects in the absence of clear data *in vivo*.

**Figure 1:**
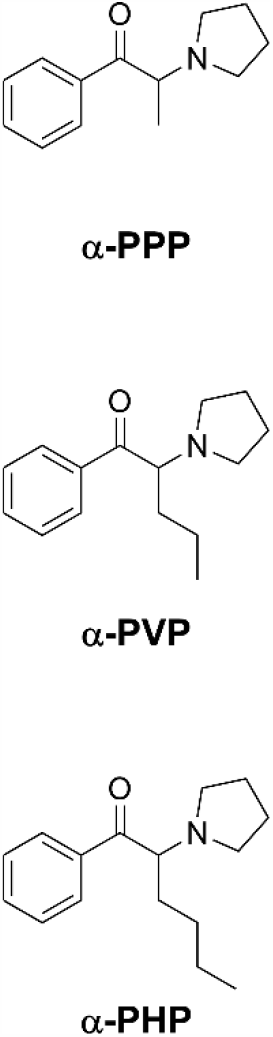
Structures of α-PPP, α-PVP and α-PHP.

Assessing behavioral effects of drugs via suppression of food motivated responding is a common behavioral pharmacological approach (Butelman et al., 1996; Jarbe et al., 2003; Lukas et al., 1988), but direct appetitive effects of specific drugs can complicate interpretation. Thus, for this study wheel running was selected as the high rate spontaneous behavior in the rat; it has the further advantage of facilitating evaluation of increased, as well as decreased, activity that may be caused by drugs (Gilpin et al., 2011; Huang, P.-K. et al., 2012; Miller et al., 2013b). We have shown that inhalation of vaporized methamphetamine and vaporized MDPV can inhibit wheel running behavior under the same conditions that it increases spontaneous locomotion (Nguyen et al., 2016; Nguyen et al., 2017). The inhalation of methamphetamine may produce enhanced behavioral effects, at a given plasma concentration, compared with even i.v. methamphetamine (Cook et al., 1993), thus, as an initial validation and a bridge of those results to the present study, we first evaluated the impact of inhaled methamphetamine, α-PVP and α-PHP on wheel activity.

Evaluation of the time course of effects of novel or emerging substances of abuse is critical. For example, a moderate dose of methamphetamine produces a rise – fall temporal pattern of change in spontaneous locomotion, whereas higher doses initially induce alternate, stereotyped behavior that actually decreases locomotion (Segal and Kuczenski, 1999). As time passes and methamphetamine levels decrease due to metabolism and excretion, spontaneous locomotor behavior rebounds, often in excess of what is observed at the same time points after saline injection. We have found such biphasic effects in the impact of *inhaled* methamphetamine on wheel activity as well (Nguyen et al., 2016). In addition, sometimes behavioral profiles can be modified by the generation of active metabolites of some compounds, which can prolong a given behavior or physiological process. Thus, full evaluation of acute effects requires approaches which are sensitive to the factor of time after injection.

Body temperature responses to restricted transporter inhibitors such as MDPV or α-PVP (Aarde et al., 2015a; Aarde et al., 2013; Kiyatkin et al., 2015; Nelson et al., 2017) are less robust in comparison with monoamine substrate/releasers such as methamphetamine or MDMA (Brown and Kiyatkin, 2004; Crean et al., 2007; Dafters and Lynch, 1998; Haughey et al., 2000; Malberg and Seiden, 1998; Miller et al., 2013a), but this is nevertheless a measure that can be used to contrast relative potency and duration of action of different drugs. By using a radiotelemetry system to evaluate body temperature, we were also able to monitor spontaneous locomotor behavior as a contrast with any observed effects on wheel running.

## 2. Materials and Methods

### 2.1 Subjects

Female (N=40) and male (N=8) Wistar rats (Charles River, New York) entered the laboratory at ∼10 weeks of age and were housed in humidity and temperature-controlled (23±1 °C) vivaria on reversed 12:12 hour light:dark cycles. All experiments were conducted in the animals’ dark (active) phase. Animals had ad libitum access to food and water in their home cages. All experimental procedures were conducted under protocols approved by the Institutional Care and Use Committee of The Scripps Research Institute in a manner consistent with the Guide for the Care and Use of Laboratory Animals (National Research Council (U.S.). Committee for the Update of the Guide for the Care and Use of Laboratory Animals. et al., 2011).

### 2.2 Drugs

The α-pyrrolidinopentiophenone HCl (α-PVP), α-pyrrolidinohexiophenone HCl (α-PHP), pentylone HCl and pentedrone HCl were obtained from Cayman Chemical. The α-pyrrolidinopropiophenone HCl (α-PPP) was obtained from Kenner Rice at the NIDA intramural research program. Methamphetamine was provided by NIDA Drug Supply. Drugs were dissolved in physiological saline for the i.p. route of administration and in propylene glycol (PG) for vapor inhalation as previously described (Nguyen et al., 2016). Doses are expressed as the salt.

### 2.3 Inhalation Apparatus

Sealed exposure chambers were modified from the 259 mm X 234 mm X 209 mm Allentown, Inc (Allentown, NJ) rat cage to regulate airflow and the delivery of vaporized drug to rats as has been previously described (Nguyen et al, 2016a; Nguyen et al, 2016b). An e-vape controller (Model SSV-1, set to 5 watts; La Jolla Alcohol Research, Inc, La Jolla, CA, USA) was triggered to deliver the scheduled series of puffs (4 10 second puffs, 2 second intervals, every 5 minutes) from Protank 3 Atomizer (Kanger Tech; Shenzhen Kanger Technology Co.,LTD; Fuyong Town, Shenzhen, China) e-cigarette cartridges by MedPC IV software (Med Associates, St. Albans, VT USA). The chamber air was vacuum-controlled by a chamber exhaust valve (i.e., a “pull” system) to flow room ambient air through an intake valve at ∼1 L per minute. This also functioned to ensure that vapor entered the chamber on each device triggering event. The vapor stream was integrated with the ambient air stream once triggered. Dosing for this study was manipulated by altering the amount of the drug in the PG solution and is expressed as mg of drug per mL of PG.

### 2.4 Wheel Apparatus

Experimental sessions were conducted in dark procedure rooms with activity wheels that attached to a typical housing chamber with a door cut to provide access to the wheel (Med Associates; Model ENV-046), using approaches previously described (Huang, P.K. et al., 2012; Nguyen et al., 2016). Rats were given access to the wheel in acute sessions during which wheel rotation (quarter-rotation resolution) activity was recorded using MED-PC IV software (Med Associates). Counts of quarter wheel rotations within a session were collected into sequential 5-min bins for initial analysis and summed into 30-minute bins for inferential statistical analysis.

### 2.5 Wheel Activity Experiments

Rats were given access to the activity wheel in a procedure room separate from the vivarium for 4 h after drug injection, or after 30 min of exposure to PG or drug concentrations infused vapor. At least one habituation session was conducted prior to the initiation of drug studies and a 3-4 day minimum interval was interposed between all active-drug test sessions.

#### 2.5.1 Female rats

A group (N=16) of female Wistar rats were initially habituated to the wheel apparatus in a single session. They were thereafter subdivided into two groups of 8 and evaluated for wheel activity immediately after 30 minutes of inhalation of either pentylone (0, 12.5, 50, 100 mg/mL in the PG) or α-PVP (0, 12.5, 50, 100 mg/mL in the PG). Thereafter, all 16 rats were evaluated for activity after 30 minutes of inhalation of methamphetamine (0, 6.25, 12.5, 100 mg/mL in the PG) vapor. Two rats died after a session, and one had to be euthanized post-session, (all a 100 mg/mL session) thus N=13 for the remaining studies. The rats were next evaluated after injection with doses of methamphetamine (0, 0.25, 1.0, 5.0 mg/kg, i.p.) in a counterbalanced order. Next, the rats were evaluated for wheel activity following injection with doses of α-PVP (0.25, 1.0, 5.0 mg/kg, i.p.) or α-PHP (0.25, 1.0, 5.0 mg/kg, i.p.) and α-PPP (0.0, 5.0, 10.0 mg/kg, i.p.). The dosing order for α-PVP and α-PHP were counterbalanced across drug by N=6/7 cohort (i.e., one cohort completed the α-PVP series first, and α-PHP series second and the other cohort completed in the opposite order) and across dose within each cohort. The highest dose of α-PHP (10 mg/kg, i.p.) was evaluated after this on a single day in all animals. The rats were then evaluated for responses to α-PPP with the doses counterbalanced within the entire N=13 sample. A 3-4 day minimum interval was interposed between all active drug test sessions.

#### 2.5.2 Male rats

A group (N=8) of male Wistar rats were habituated the wheel apparatus in two sessions, the first with no prior treatment and the second with a saline injection (i.p.). Thereafter, the group was assessed for wheel activity after injection with doses of α-PVP (0, 0.25, 1.0, 5.0 mg/kg, i.p.) or α-PHP (0, 0.25, 1.0, 5.0 mg/kg, i.p.) with the dosing order counterbalanced both within and across compounds. One animal was found dead overnight due to unknown causes during this series of studies, thus N=7 for the analysis and subsequent studies. Next the rats were evaluated for wheel activity after injection with vehicle or 10 mg/kg, i.p., of α-PPP, α-PVP or α-PHP in a counterbalanced order.

### 2.6 Radiotelemetry

#### 2.6.1 Surgical implantation

A group of female (N=8) Wistar rats were implanted with sterile radiotelemetry transmitters (Data Sciences International, St Paul, MN) in the abdominal cavity under isoflurane/oxygen inhalation anesthesia (isoflurane 5% induction, 1–3% maintenance) using sterile procedures, as previously described (Taffe et al., 2015; Wright et al., 2012). For the first 3 days of the recovery period, an antibiotic Cefazolin (Hikma Farmaceutica, Portugal; 0.4 mg/kg, i.m. in sterile water day 1, s.c. days 2–3) and an analgesic flunixin (FlunixiJect, Bimeda USA, Oakbrook Terrace, IL; 2.5 mg/kg, s.c. in saline) were administered daily. Following surgical recovery (minimum of 7 days), animals were evaluated in clean standard plastic home cages (thin layer of bedding) in a dark testing room, separate from the vivarium, during the (vivarium) dark cycle. Radiotelemetry transmissions were collected via telemetry receiver plates (Data Sciences International, St Paul, MN; RPC-1 or RMC-1) placed under the cages with data collected as body temperature and activity rate (counts per minute) every 5 minutes.

#### 2.6.2 Experimental Procedure

Test sessions for inhalation or injection studies started with at least a 30-minute baseline interval followed by vapor exposure or injection of the scheduled drug / vehicle doses. These individuals contributed to similar studies involving the injection of pentylone, pentedrone and methylone, as was described in previously published investigations (Javadi-Paydar et al., 2018b). The timeline for studies for this group was α-PPP (0, 1.0, 5.0, 10.0 mg/kg, i.p.), reported herein, the three previously published drug challenges, then α-PHP (0, 1.0, 5.0, 10.0 mg/kg, i.p.) and finally α-PVP (0, 1.0, 5.0, 10.0 mg/kg, i.p.). Approximately 60 days elapsed between the final α-PPP challenge day and the first α-PHP challenge day, thus a repetition of the 10 mg/kg α-PPP dose was conducted after completion of the α-PVP studies. In all studies other than this last for these animals, the drug doses were evaluated in a counter-balanced order within drug identity, with no more than 2 active drug doses experienced in a week (i.e., a 3-4 day inter-drug interval).

### 2.7 Intracranial Self-Stimulation (ICSS) Reward Procedure

For these studies, procedures were adapted from well-established protocols described previously (Kenny et al., 2006; Kornetsky et al., 1979; Markou and Koob, 1992; Nguyen et al., 2016). Rats (N=16) were surgically prepared with unilateral electrodes aimed at the medial forebrain bundle (coordinates: AP −0.5mm, ML ±1.7mm, DV skull −9.5mm) under isoflurane/oxygen inhalation anesthesia (isoflurane 5% induction, 1–3% maintenance) using sterile techniques. For the first 3 days of the recovery period, an antibiotic Cefazolin (Hikma Farmaceutica, Portugal; 0.4 mg/kg, i.m. in sterile water day 1, s.c. days 2–3) and an analgesic flunixin (FlunixiJect, Bimeda USA, Oakbrook Terrace, IL; 2.5 mg/kg, s.c. in saline) were administered daily. Following surgical recovery (minimum 14 days) rats were trained in a procedure adapted from the discrete-trial current-threshold procedure as described in (Nguyen et al., 2016). After rats had been trained to stability, challenge studies commenced wherein vehicle or doses of test drugs were administered before the session as follows. Two individuals did not pass the training phase and two had their ports chewed beyond repair by a cage mate, all individuals were subsequently single-housed to prevent further port damage. All injections were conducted 15 minutes prior to the start of the ICSS session. In the first experiment, rats (N=11 completed all conditions) were injected with saline or MA (0.5 mg/kg, i.p.) in a counterbalanced order across the group. MA was included as a positive control, since this reliably reduces thresholds in male rats in our hands (Nguyen et al., 2016; Nguyen et al., 2019), similar to the effects of 3,4-methylenedioxymethamphetamine (Aarde et al., 2017). The rats (N=10 completed all conditions) were next injected 15 minutes prior to the session with saline or one of two doses (0.5, 1.0 mg/kg, i.p.) of α-PVP or α-PPP in a counter-balanced order. In a subsequent experiment, rats were injected 15 minutes prior to the session with saline or one of two doses (0.5, 1.0 mg/kg, i.p.) of pentylone or pentedrone in a counter-balanced order.

### 2.8 Data Analysis

Data (wheel quarter rotations, body temperature and spontaneous activity rates) were analyzed by ANOVA with repeated measures factors for dose and time post-injection or time after the end of inhalation for vapor studies. A mixed-effects model was substituted whenever there were data missing for an analysis cell. The wheel-activity and telemetry data were collected on a 5-minute schedule but are expressed as 30-minute averages for analysis and presentation. Any missing temperature values were interpolated from the values before and after the lost time point. Activity rate values were not interpolated because 5-minute to 5-minute values can change dramatically, thus there is no justification for interpolating. ICSS thresholds were represented as a percent of the individual threshold value obtained on the day before the challenge day. A criterion of P<0.05 was used to infer that a significant difference existed. Any significant main effects on temperature and activity were followed with post-hoc analysis using Tukey (multi-level factors) or Sidak (two-level factors) correction. Significant main effects of drug treatments on ICSS thresholds were assessed using the Dunnett procedure to compare with saline injection. All analysis used Prism for Windows (v. 8.3; GraphPad Software, Inc, San Diego CA).

## 3. Results

### 3.1 Suppression of wheel activity in female rats

Female rats engage in significant wheel activity when presented with the wheel for 4 h intermittently, i.e., ∼2 times per week. Under baseline (not shown) and vehicle injection conditions, the rats express the most activity in the first 30 minutes, and roughly stable levels from 60-150 minutes. Activity levels sometimes decline in the final 90 minutes of the session. The initial, positive control studies examined wheel activity after methamphetamine (0.25, 0.5, 5 mg/kg, i.p.) injection and methamphetamine vapor (6.25, 12.5, 100 mg/mL in PG) inhalation for 30 minutes (**Figure 2**). Only 6 rats completed the 6.25 mg/mL condition due to experimental exigencies.

**Figure 2:**
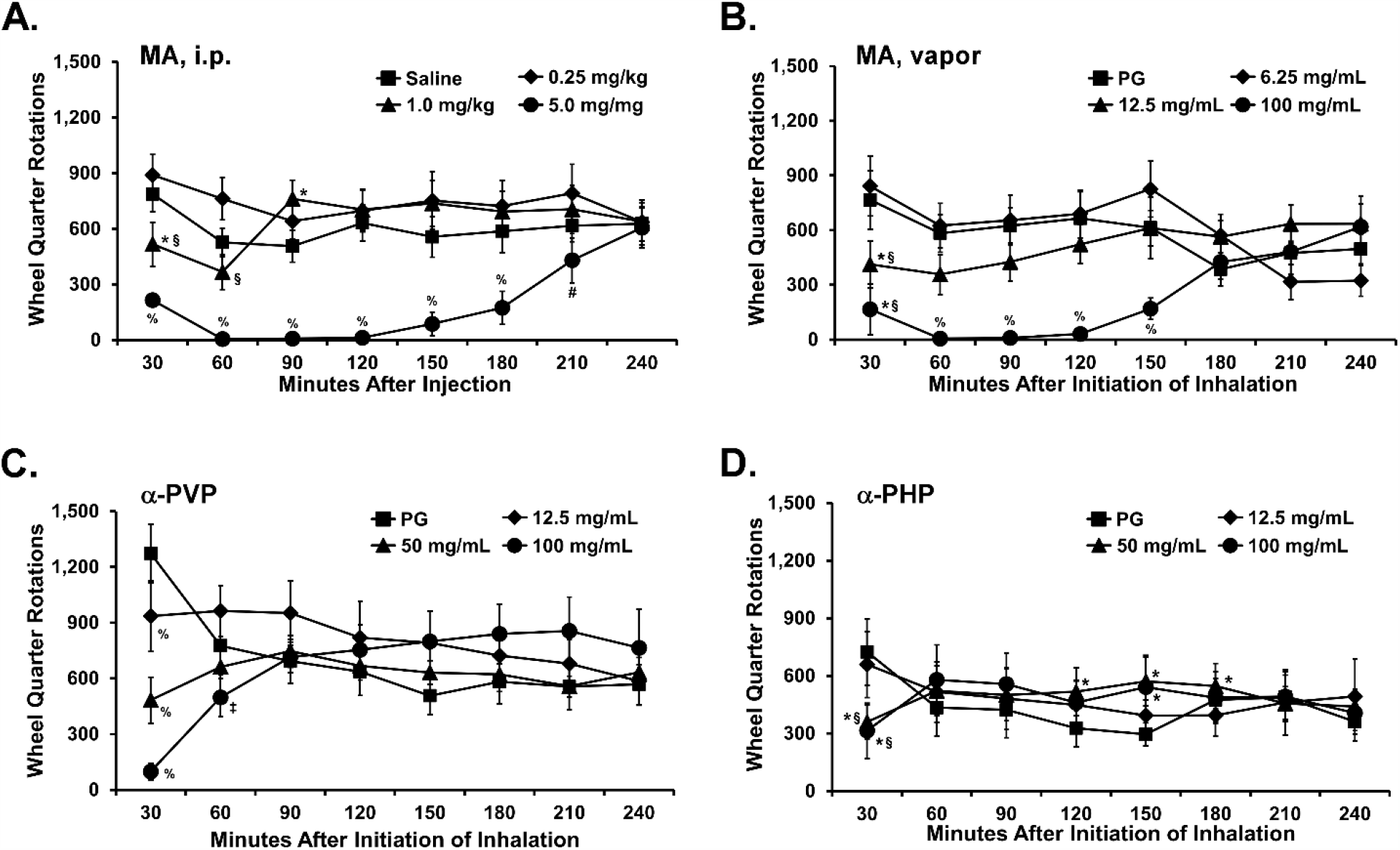
Mean Wheel activity after A) methamphetamine injection (N=13), B) methamphetamine vapor inhalation (N=13 except N=6 for the 6.25 mg/mL condition), C) α-PVP (N=8) vapor inhalation and D) α-PHP (N=13) vapor inhalation; A significant difference from Saline or PG is indicated with *, from the lowest dose with §, from the intermediate dose with ‡, from the lowest and intermediate doses with #, and a significant difference from all other dosing conditions, within drug, is indicated with %.

Methamphetamine vapor inhalation, or injection, at the highest doses (100 mg/mL and 5 mg/kg, respectively) produced nearly a complete suppression of wheel activity for 2 hours after injection or after the end of inhalation (**Figure 2 A**,**B**). No dose produced a sustained increase in wheel activity. For the i.p. injection study, the ANOVA confirmed a significant effect of time after vapor initiation [F (7, 84) = 4.29; P<0.0005], of dose condition [F (3, 36) = 14.69; P<0.0001] and the interaction of dose condition with time after vapor initiation [F (21, 252) = 4.42; P<0.0001] on wheel activity. The post-hoc test further confirmed that activity was significantly lower 30-180 minutes after injection of the 5.0 mg/kg methamphetamine dose, compared with injection of saline and either lower dose. After injection of 1.0 mg/kg, activity was lower compared with saline 30-minutes after injection but significantly higher at the 90 minute time-point.

For the methamphetamine vapor inhalation study, the mixed-effects analysis confirmed a significant effect of dose condition [F (3, 39) = 7.48; P<0.0005] and the interaction of dose condition with time after vapor cessation [F (21, 185) = 4.69; P<0.0001] on wheel activity (**Figure 2 B**). The post-hoc test further confirmed that activity was lower in the 30-150 minutes after the end of the 100 mg/mL inhalation compared with either PG or 6.25 mg/mL conditions and 60-150 minutes after the end of inhalation compared with the 12.5 mg/mL condition. Wheel activity was also significantly lower 30 minutes after the end of inhalation of 12.5 mg/mL compared with after PG or 6.25 mg/mL inhalation.

Inhalation of α-PVP and α-PHP vapor also produced a dose dependent reduction of wheel activity however this lasted only for the first 30 minutes of wheel access (**Figure 2 C, D**). The ANOVAs confirmed a significant effect of the interaction of vapor inhalation dose with time after vapor cessation for α-PVP (F (21, 147) = 6.22; P<0.0001) and α-PHP (F (21, 252) = 4.37; P<0.0001); there was no significant effect of either factor alone for either drug. The post-hoc test confirmed that wheel activity was significantly lower in the initial 30 minutes after the inhalation of 50 or 100 mg/mL of α-PVP compared with the PG or 12.5 mg/mL conditions. Wheel activity also differed significantly between the 50 and 100 mg/mL conditions, as well as between the PG and 12.5 mg/mL conditions in the initial 30-minute time-bin.

Finally, wheel activity differed significantly 60 minutes after the cessation of inhalation of 100 mg/mL α-PVP compared with the 12.5 mg/mL condition. The post-hoc test further confirmed that wheel activity was significantly lower in the initial 30 minutes after inhalation of 50 or 100 mg/mL of α-PHP compared with the PG or 12.5 mg/mL conditions. Conversely, wheel activity was significantly *higher*, compared with after PG inhalation, following the 100 mg/mL (150 minutes after the end of inhalation) or 50 mg/mL (120-180 minutes after the end of inhalation) α-PHP conditions.

Intraperitoneal injection of α-PPP, α-PVP and α-PHP also reduced wheel activity in the female rats (**Figure 3**). There was a dose-related suppression of wheel activity after injection of α-PPP and the ANOVA confirmed significant effects of Time post-injection [F (7, 84) = 10.36;P<0.0001], of Dose [F (2, 24) = 4.26, P<0.05] and of the interaction of factors [F (14, 168) = 2.99; P<0.0005] on wheel activity. Likewise, the ANOVA for the α-PVP experiment confirmed significant effects of Dose [F (3, 36) = 3.13; P<0.05] and of the interaction of Dose with Time post-injection [F (21, 252) = 2.47; P<0.001] on wheel activity. Finally, the ANOVA for the α-PHP investigation confirmed significant effects of Time post-injection [F (7, 84) = 3.79; P<0.005] and of the interaction of factors [F (28, 336) = 4.8; P<0.0001] on wheel activity.

**Figure 3:**
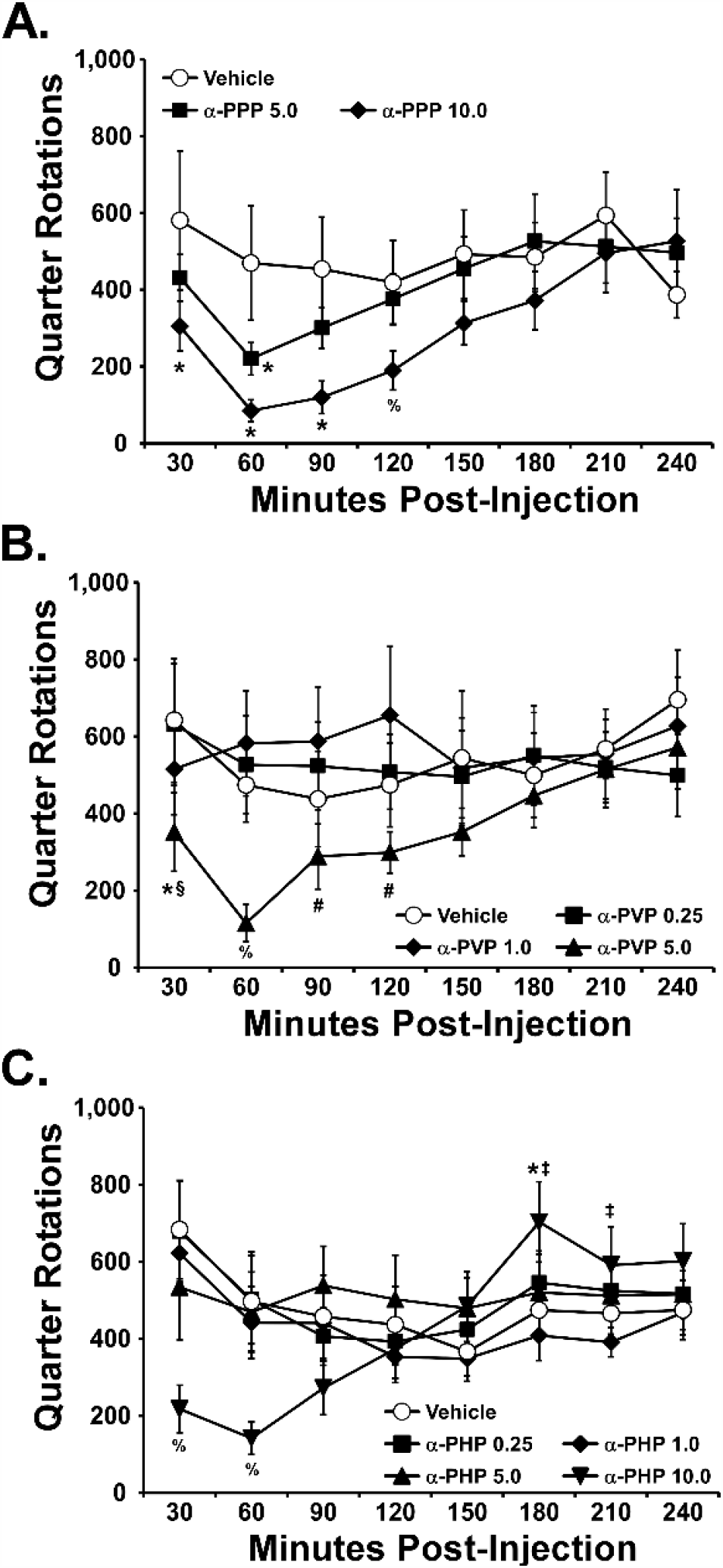
Mean wheel activity of female rats (n=8; ±SEM) after injection with doses of A) α-PPP, B) α-PVP, or C) α-PHP. A significant difference from vehicle at a given time post-injection is indicated with *, from the 0.25 mg/kg dose with §, from the 1.0 mg/kg dose with ‡, from the 0.25 and 1.0 mg/kg doses with #, and a significant difference from all other dosing conditions, within drug, is indicated with %.

### 3.2 Suppression of wheel activity in male rats

Male rats also engaged in significant wheel activity when presented with the wheel for 4 h intermittently (∼2 times per week) and, under vehicle injection condition, express the most activity in the first 30 minutes (**Figure 4**). Unlike the female rats, while roughly stable levels of activity were expressed for the 60-150 minute interval after the wheel was made accessible, activity declined significantly in the last 90 minutes of the session. Injection of α-PVP and α-PHP produced dose-related alterations of wheel activity in male rats, with significant suppression observed in the initial hour after the 10 mg/kg dose. The ANOVAs confirmed significant effects of Time, Dose and/or the Interaction of factors on activity for the α-PVP [Dose: F (4, 24) = 7.84; P<0.0005; Time: F (7, 42) = 10.79; P<0.0001; Interaction: F (28, 168) = 1.27; P=0.1812], α-PHP [F (4, 24) = 5.83; P<0.005; Time: F (7, 42) = 5.36; P<0.0005; Interaction: F (28, 168) = 2.94; P<0.0001], and 10 mg/kg contrast [Dose: F (3, 18) = 6.68; P<0.005; Time: F (7, 42) = 3.28; P<0.01; Interaction: F (21, 126) = 2.22; P<0.005] studies. The post-hoc test of the marginal means for drug treatment condition confirmed that activity was lower after 10 mg/kg α-PVP dose compared with after the 1 or 5 mg/kg α-PVP doses (**Figure 4 A**). Wheel activity was significantly higher after the 5 mg/kg α-PHP dose compared with each of the other treatment conditions (**Figure 4 B**). Finally, in the 10 mg/kg comparison, activity was significantly lower after α-PVP compared with either α-PPP or α-PHP (**Figure 4 C**).

**Figure 4:**
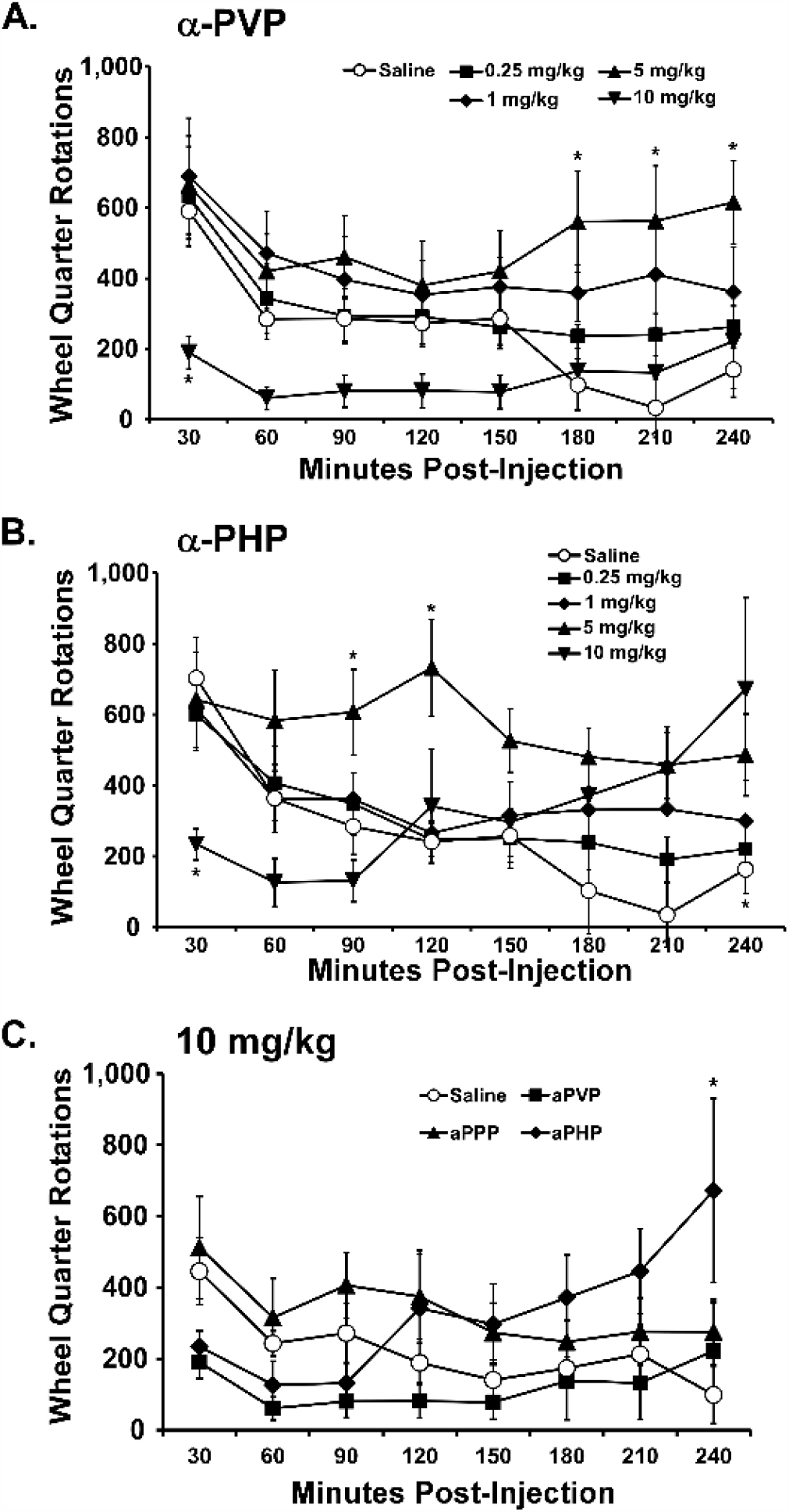
Mean wheel activity of male rats (n=7; SEM) after injection of A) α-PVP or B) α-PHP. C) A direct contrast of the effects of injection of 10 mg/kg of α-PPP with the results for 10 mg/kg α-PVP and α-PHP. A significant difference from vehicle at a given time post-injection is indicated with *.

### 3.5 Activity and body temperature

#### 3.5.1 Original Dose-Response

Injection of all three drugs altered the activity and body temperature of female rats (**Figure 5**). The ANOVAs confirmed significant effects of Time, Dose and/or the interaction of factors on <u>activity</u> for the α-PPP [Dose: F (3, 21) = 15.28, P<0.0001; Time: F (8, 56) = 23.75, P<0.0001; Interaction: F (24, 168) = 4.49, P<0.0001], α-PVP [Dose: F (3, 21) = 1.73, P=0.1924; Time: F (8, 56) = 2.43, P<0.05; Interaction: F (24, 168) = 2.48, P<0.0005], and α-PHP [Dose: F (3, 21) = 8.7, P<0.001; Time: F (8, 56) = 3.73, P<0.005; Interaction: F (24, 168) = 5.42, P<0.0001], studies. The ANOVAs also confirmed significant effects of Time, Dose and/or the interaction of factors on body <u>temperature</u> for the α-PPP [Dose: F (3, 21) = 2.50, P=0.0871; Time: F (8, 56) = 18.95, P<0.0001; Interaction: F (24, 168) = 4.40, P<0.0001], α-PVP [Dose: F (3, 21) = 3.54, P<0.05; Time: F (8, 56) = 6.91, P<0.0001; Interaction: F (24, 168) = 3.12, P<0.0001] and α-PHP [Dose: F (3, 21) = 2.37, P=0.0992; Time: F (8, 56) = 6.33, P<0.0001; Interaction: F (24, 168) = 4.16; P<0.0001] studies. The post-hoc test confirmed that body temperature was elevated by α-PPP and decreased by α-PVP and α-PHP.

**Figure 5:**
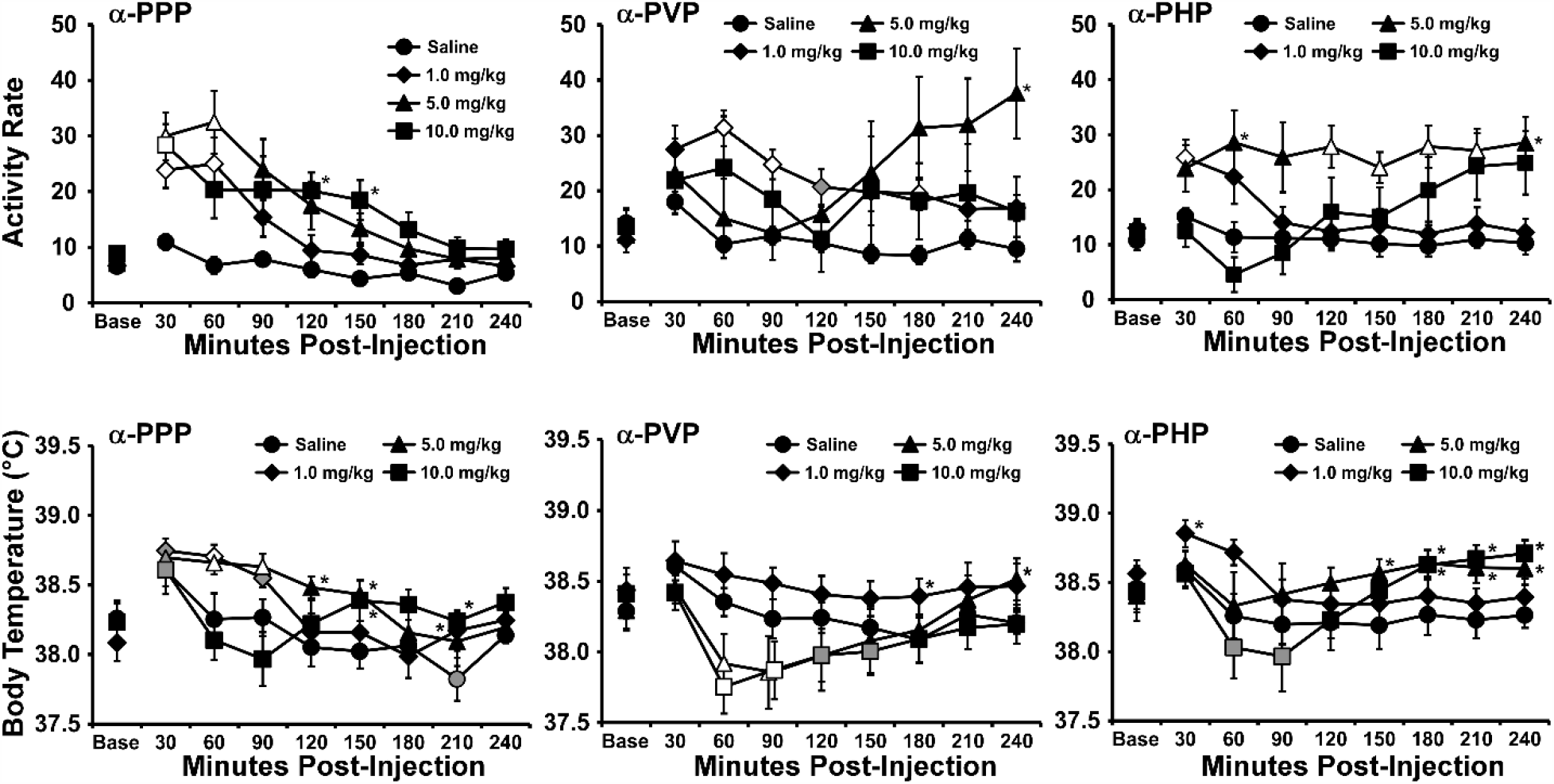
Mean (N=8; ±SEM) activity responses of female Wistar rats after injection with doses (1, 5, 10 mg/kg, i.p.) of α-PPP, α-PVP or α-PHP. A difference from vehicle (but not baseline) is indicated with *.

#### 3.5.2. Repetition of α-PPP

This analysis confirmed that the profile of response to 10 mg/kg α-PPP at Time 2 was more similar to the response to the 10 mg/kg dose of α-PHP or α-PVP, and less similar to the effects of the same α-PPP at Time 1 (**Figure 6**). Whether due to the age of the animals or a degree of sensitization due to the repeated studies in the group, it indicates that α-PPP is indeed similarly potent as the other two analogs when compared directly. The statistical analysis confirmed effects of Time after injection [F (2.319, 16.23) = 5.23; P<0.05], of treatment condition [F (1.721, 12.04) = 9.64; P<0.005] and of the interaction of factors [F (3.718, 26.03) = 5.13; P<0.005] on activity rate. The post-hoc test confirmed that activity was higher than the baseline and respective vehicle condition 60 minutes after injection of α-PPP at Time 1 but not Time 2. Activity was also significantly higher 180-210 minutes after injection of α-PPP at Time 2 compared with the effects of α-PPP at Time 1. There was also a small but statistically reliable reduction of body temperature at Time 2, similar to what was observed for α-PHP and α-PVP (**Figure 5**). The statistical analysis confirmed effects of Time after injection [F (2.582, 18.08) = 13.06; P<0.0005], and of the interaction of factors [F (3.832, 26.82) = 5.78; P<0.005] on body temperature. The post-hoc test confirmed temperature was lower than the baseline, and the respective vehicle condition, 60 min after injection of α-PPP at Time 2.

**Figure 6:**
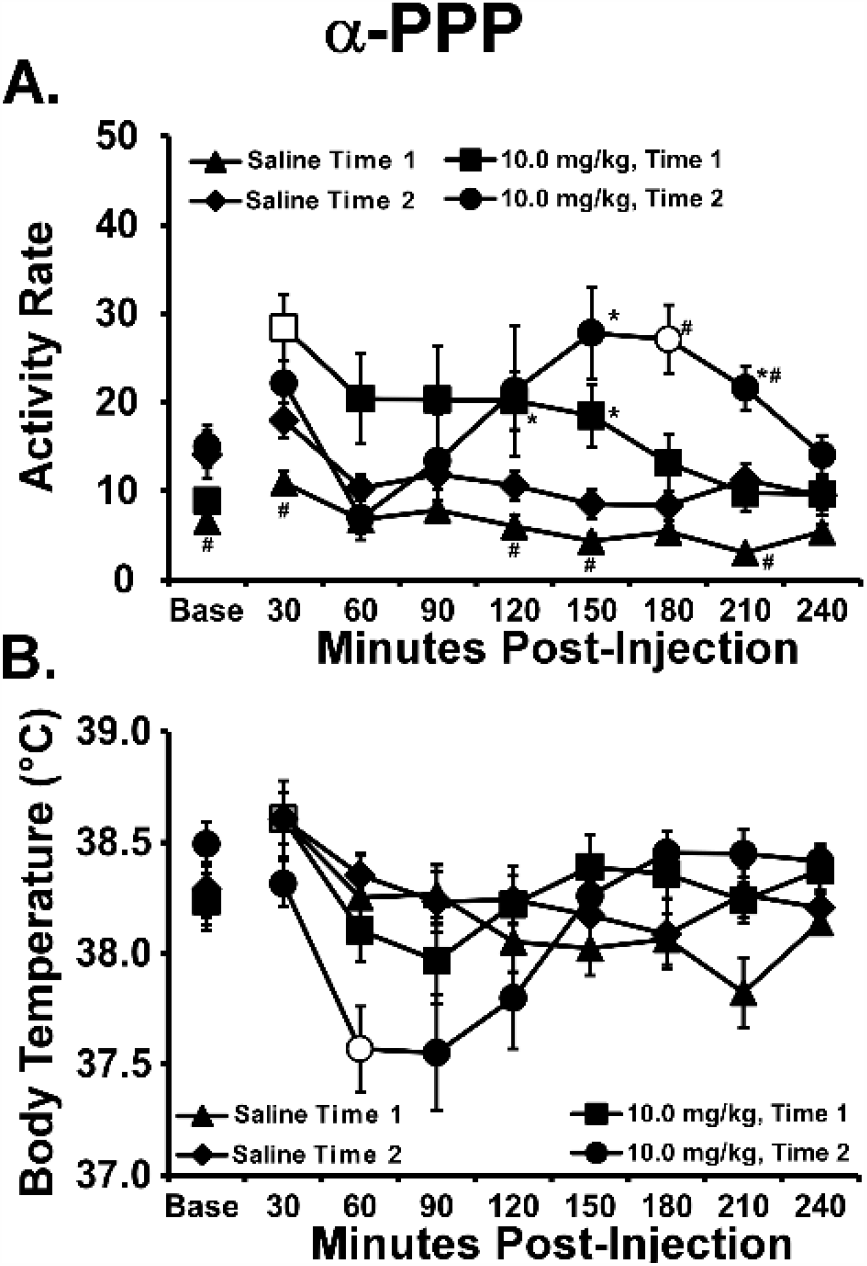
Mean (N=8; ±SEM) A) Activity rate and B) body temperature responses of female rats to injection of α-PPP (10 mg/kg, i.p.) or vehicle on Time 1 (the original study) and Time 2 (a replication of this dose after completion of the α-PVP and α-PHP studies). Shaded symbols indicate a significant difference from the Baseline (Base), within treatment condition. Open symbols indicate a significant difference from the Baseline, within treatment condition and from the Vehicle condition, within Time. A significant difference from Time 1 to Time 2 within drug treatment is indicated with # and a difference from the vehicle (but not baseline) within Time is indicated with *.

### 3.6 Brain Reward Threshold

The injection of α-PPP and α-PVP lowered reward thresholds in female (N=11) Wistar rats (**Figure 7**) as was confirmed by a significant main effect of drug pre-treatment [F (3.862, 38.62) = 5.90; P<0.001] on reward thresholds. The Dunnett post-hoc test first confirmed the positive control validation, since 0.5 mg/kg methamphetamine significantly lowered thresholds compared with the first vehicle injection. More limited, but significant reductions in reward threshold were produced by α-PVP (0.5 mg/kg, i.p.) and α-PPP (1.0 mg/kg, i.p.). There was likewise a significant effect of pre-treatment condition in the second study (N=10) as confirmed by the ANOVA [F (2.421, 21.79) = 5.69; P<0.01]. The Dunnett post-hoc test confirmed that thresholds were lower than after saline injection when pentylone (0.5 mg/kg, i.p.) or pentedrone (1.0 mg/kg, i.p.) was administered.

**Figure 7:**
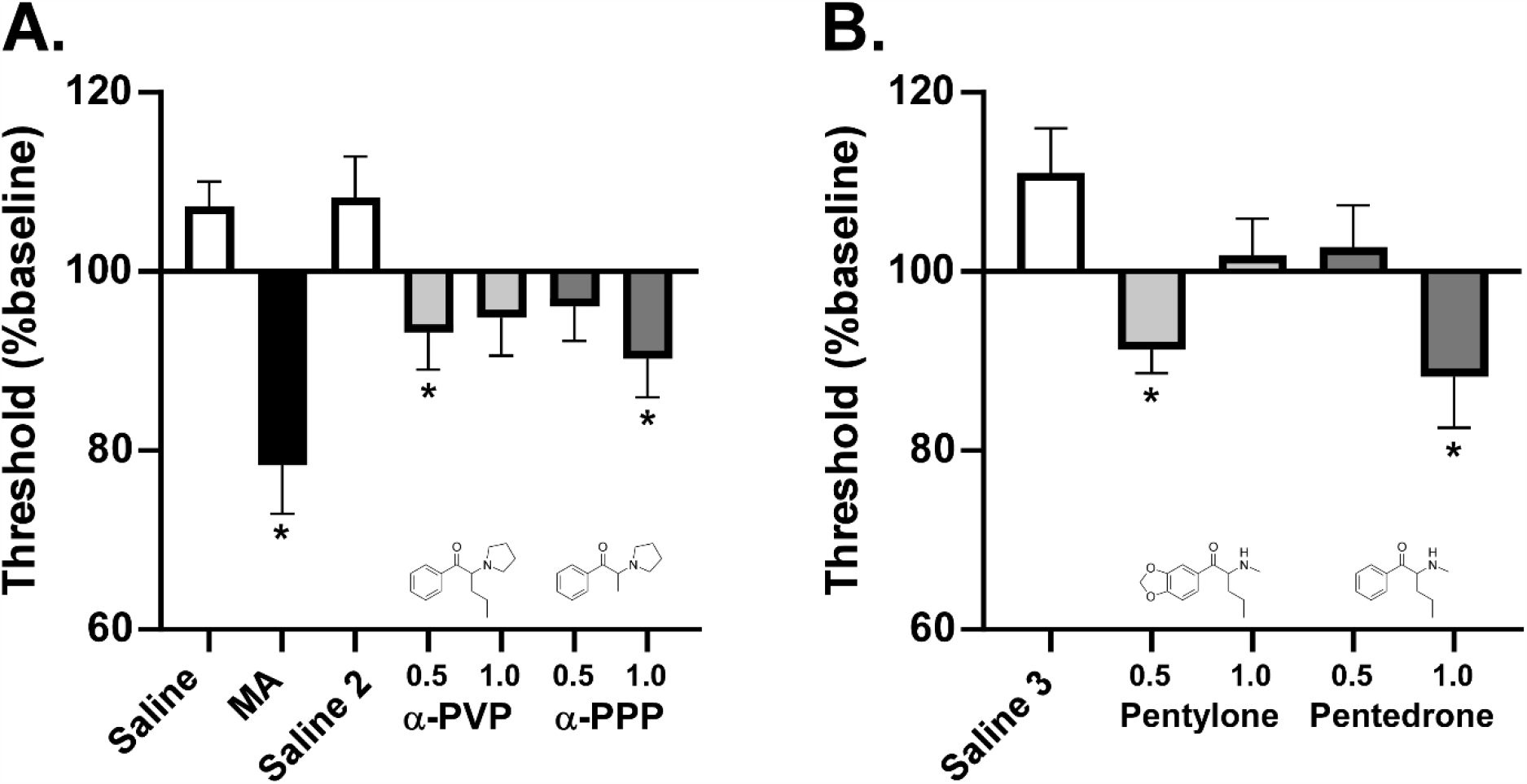
Female Wistar rats (N=11) injected with saline or methamphetamine (MA; 0.5 mg/kg, i.p. 15 minutes before session), α-PVP and α-PPP (0.5, 1.0 mg/kg, i.p.). Female Wistar rats (N=10) injected with saline or pentylone or pentedrone (0.5, 1.0 mg/kg, i.p.). A significant difference from Saline and Saline 3, respectively, is indicated with *.

## 4. Discussion

This investigation demonstrates that cathinone derivatives α-pyrrolidinopentiophenone (α-PVP), α-pyrrolidinohexiophenone (α-PHP) and α-pyrrolidinopropiophenone (α-PPP) produce similar psychomotor stimulant effects in the rat. First, spontaneous locomotor activity within the normal home cage environment was increased for hours by each drug. That is, following injection, spontaneous activity was increased and then declined approximately monotonically with time at moderate doses (**Figure 5**). Second, higher doses interfered with the expression of an otherwise high-rate behavior, i.e. wheel activity in a dose- and time-dependent manner (**Figures 3**,**4**). This occurred for the α-PVP and α-PHP analogs in the male rats and all three analogs in the female rats. Thirdly, body temperature was increased at lower doses, and decreased by higher doses, of all three drugs (**Figure 5, 6**). Finally, α-PVP and α-PPP reduced reward thresholds in the intracranial self-stimulation (ICSS) procedure (**Figure 7**), consistent with pro-reward effects shown for other psychomotor stimulants such as methamphetamine (Nguyen et al., 2016).

Although we have shown previously that the inhalation of MDPV using an Electronic Drug Delivery System (EDDS; aka, “e-cigarette”) can alter locomotor activity and brain reward thresholds in rats (Nguyen et al., 2016), this is also the first demonstration that the α-pyrrolidino-phenone compounds can also be effectively administered by vapor inhalation (**Figure 2**). This lends credence to the still poorly confirmed allegations in popular media that this route is used for α-PVP self-administration by some humans (Anderson, 2015; D’Oench, 2015; Milian, 2015). The relatively short duration of action of α-PVP and α-PHP in suppressing wheel activity, compared with the duration of effects after injection, further confirms the additional risks of the inhalation route. That is, a rapid offset after an approximately similar maximum effect (as in Cook et al. 1993) is likely to induce frequent re-dosing during an acute use episode. This may further exacerbate a binge-like initial acquisition phenotype similar to that which has been observed in rats under some intravenous self-administration conditions for α-PVP and MDPV (Aarde et al., 2015b; Javadi-Paydar et al., 2018a).

The α-PPP compound is less potent than the other two since it produced no impact on wheel activity in male rats at a dose (10 mg/kg, i.p.) where the α-PHP and α-PVP analogs robustly suppressed initial wheel activity. It was also less potent compared with α-PVP in the ICSS procedure in female rats. This is consistent with prior demonstrations of reduced potency of α-PPP in intravenous self-administration (Gannon et al., 2018; Nagy et al., 2020) as well as in assays of dopamine transporter inhibition (Gannon et al., 2018; Kolanos et al., 2015). Nevertheless, α-PPP is clearly an effective stimulant since it reduced wheel running and increased spontaneous locomotor behavior in female rats. More subtle differences emerged between the α-PHP and α-PVP analogs as the former appeared to permit rebound wheel activity ∼3-4 hours after injection whereas activity only approximated that of the saline condition 3-4 hours after α-PVP injection at the same dose. In addition, wheel activity was actually higher than that exhibited during saline ∼2 hours after 5 mg/kg α-PHP but only in the 3-4 hour interval after 5 mg/kg α-PVP. This pattern suggests a more rapid metabolism of α-PHP compared with α-PVP.

The temperature response to all three drugs was small (**Figure 5, 6**), consistent with prior reports for α-PVP and MDPV (Aarde et al., 2015a; Aarde et al., 2013), as well as for other non-pyrrolidino-phenone cathinones with mostly monoamine transporter inhibitor activity (Javadi-Paydar et al., 2018b). This stands in substantial contrast with the larger temperature effects produced by some of the monoamine transporter substate/releaser cathinones (Miller et al., 2013a; Wright et al., 2012) and the substrate/releaser amphetamines (Aarde et al., 2017; Malberg and Seiden, 1998; Miller et al., 2013a). These results contrast slightly with one prior report of thermoregulatory effects of α-PVP in male and female Sprague-Dawley rats, however those studies were conducted in the light part of the cycle and relative elevations of temperature were most pronounced several hours after injection when temperature declined in the vehicle condition (Nelson et al., 2019). A similar caveat applies to the locomotor data in that study, where the inference of increased locomotion was driven by a decrease in activity after vehicle injection rather than any change from the baseline in the active drug conditions.

The intracranial self-stimulation reward paradigm used here consistently shows reduced thresholds following injection of amphetamine (Esposito et al., 1980), mephedrone, MDPV or methamphetamine (Nguyen et al., 2016) and MDMA (Aarde et al., 2017; Hubner et al., 1988). MDMA is less potent than MA in reducing thresholds (Aarde et al., 2017; Nguyen et al., 2016), but the 3,4-methylenedioxy motif is not a consistent predictor of that, as was illustrated by pentylone exhibiting higher potency relative to pentedrone in this study (**Figure 7**).

In total, this study further confirms the considerable limitations in predicting relative efficacy, *in vivo*, from either structure-activity inferences based on other compounds, or from *in vitro* pharmacology. For the compounds compared in this study, either of those might have predicted greater potency or efficacy differences between the α-PHP and α-PPP analogs than was actually observed. Consistent with this prediction, intravenous self-administration data using rats suggest differential potency, but similar efficacy of α-PVP and α-PPP, as well as of the 3,4-methylenedioxy substituted analogs of these two structures (Gannon et al., 2018). Furthermore, when considering that human users actively seek to titrate the drug dosing to achieve the intended effect, it is clear that only small differences in drug intake would be required to achieve a similar effect for these compounds, depending on which of the three analogs the person happened to obtain. As a minor caveat to the general inference of similarity that may be drawn from these specific data, it may be the case that other behavioral or physiological effects of these drugs, not studied herein, might be affected more differently by one of the compounds compared with another.

## Literature Cited

Aarde, S.M., Creehan, K.M., Vandewater, S.A., Dickerson, T.J., Taffe, M.A., 2015a. In vivo potency and efficacy of the novel cathinone alpha-pyrrolidinopentiophenone and 3,4-methylenedioxypyrovalerone: self-administration and locomotor stimulation in male rats. Psychopharmacology 232(16), 3045–3055.

Aarde, S.M., Huang, P.K., Creehan, K.M., Dickerson, T.J., Taffe, M.A., 2013. The novel recreational drug 3,4-methylenedioxypyrovalerone (MDPV) is a potent psychomotor stimulant: self-administration and locomotor activity in rats. Neuropharmacology 71, 130–140.

Aarde, S.M., Huang, P.K., Dickerson, T.J., Taffe, M.A., 2015b. Binge-like acquisition of 3,4-methylenedioxypyrovalerone (MDPV) self-administration and wheel activity in rats. Psychopharmacology 232(11), 1867–1877.

Aarde, S.M., Huang, P.K., Taffe, M.A., 2017. High ambient temperature facilitates the acquisition of 3,4-methylenedioxymethamphetamine (MDMA) self-administration. Pharmacology, biochemistry, and behavior 163, 38–49.

Adamowicz, P., Gieron, J., Gil, D., Lechowicz, W., Skulska, A., Tokarczyk, B., 2016. The prevalence of new psychoactive substances in biological material - a three-year review of casework in Poland. Drug testing and analysis 8(1), 63–70.

Anderson, C., 2015, Flakka, Synthetic Drug Behind Increasingly Bizarre Crimes. Associated Press. The Associated Press. http://hosted.ap.org/dynamic/stories/U/US_FLAKKAS_EMERGENCE

Beck, O., Backberg, M., Signell, P., Helander, A., 2018. Intoxications in the STRIDA project involving a panorama of psychostimulant pyrovalerone derivatives, MDPV copycats. Clinical toxicology (Philadelphia, Pa 56(4), 256–263.

Benzie, F., Hekman, K., Cameron, L., Wade, D.R., Smolinske, S., 2011. Emergency department visits after use of a drug sold as “bath salts” ---michigan, november 13, 2010--march 31, 2011. MMWR Morb Mortal Wkly Rep 60(19), 624–627.

Bonson, K.R., Dalton, T., Chiapperino, D., 2019. Scheduling synthetic cathinone substances under the Controlled Substances Act. Psychopharmacology 236(3), 845–860.

Brown, P.L., Kiyatkin, E.A., 2004. Brain hyperthermia induced by MDMA (ecstasy): modulation by environmental conditions. Eur J Neurosci 20(1), 51–58.

Butelman, E.R., Negus, S.S., Lewis, J.W., Woods, J.H., 1996. Clocinnamox antagonism of opioid suppression of schedule-controlled responding in rhesus monkeys. Psychopharmacology 123(4), 320–324.

Cook, C.E., Jeffcoat, A.R., Hill, J.M., Pugh, D.E., Patetta, P.K., Sadler, B.M., White, W.R., Perez-Reyes, M., 1993. Pharmacokinetics of methamphetamine self-administered to human subjects by smoking S-(+)-methamphetamine hydrochloride. Drug metabolism and disposition: the biological fate of chemicals 21(4), 717–723.

Crean, R.D., Davis, S.A., Taffe, M.A., 2007. Oral administration of (+/-)3,4-methylenedioxymethamphetamine and (+)methamphetamine alters temperature and activity in rhesus macaques. Pharmacology, biochemistry, and behavior.

D’Oench, P., 2015, Exclusive: Surveillance Tape Shows Mom Accused Of Abandoning Baby After Taking Flakka. CBSMiami. CBSMiami. http://miami.cbslocal.com/2015/05/04/exclusive-surveillance-tape-shows-mom-accused-of-abandoning-baby-after-taking-flakka/

Dafters, R.I., Lynch, E., 1998. Persistent loss of thermoregulation in the rat induced by 3,4-methylenedioxymethamphetamine (MDMA or “Ecstasy”) but not by fenfluramine. Psychopharmacology 138(2), 207–212.

DEA, 2019a. National Forensic Laboratory Information System: Drug 2018 Annual Report. U.S. Drug Enforcement Administration, Springfield, VA.

DEA, 2019b. Schedules of Controlled Substances: Temporary Placement of NEthylhexedrone, α-PHP, 4-MEAP, MPHP, PV8, and 4-Chloro-α-PVP in Schedule I, in: rdJustice, D.o. (Ed.) 84 ed. Department of Justice, Washington, DC, pp. 18423–18428.

DEA, 2020. National Forensic Laboratory Information System: Drug 2019 Midyear Report. U.S. Drug Enforcement Administration, Springfield, VA.

Drug Enforcement Administration, D.o.J., 2014. Schedules of controlled substances: temporary placement of 10 synthetic cathinones into Schedule I. Final order. Federal register 79(45), 12938–12943.

Drug Enforcement Administration, D.o.J., 2017. Schedules of Controlled Substances: Placement of 10 Synthetic Cathinones Into Schedule I. Final rule. Federal register 82(39), 12171–12177.

Eshleman, A.J., Wolfrum, K.M., Reed, J.F., Kim, S.O., Swanson, T., Johnson, R.A., Janowsky, A., 2017. Structure-Activity Relationships of Substituted Cathinones, with Transporter Binding, Uptake, and Release. The Journal of pharmacology and experimental therapeutics 360(1), 33–47.

Esposito, R.U., Perry, W., Kornetsky, C., 1980. Effects of d-amphetamine and naloxone on brain stimulation reward. Psychopharmacology 69(2), 187–191.

Fisher, D., 2019, Sullivan County officials hail effort to curb ‘bath salt’ use. The Union Leader. Union Leader Corp. https://www.unionleader.com/news/crime/sullivan-county-officials-hail-effort-to-curb-bath-salt-use/article_d4d63426-7237-53f2-9e05-2cb7a42bac5a.html

Fox12Staff, 2019, DOJ: Vancouver man arrested for trafficking ‘bath salts’ out of mobile home, storage units Fox 12 Oregon. KPTV-KPDX Broadcasting Corporation. https://www.kptv.com/news/doj-vancouver-man-arrested-for-trafficking-bath-salts-out-of/article_fb0c1642-0289-11ea-a438-4faa38889331.html

Gannon, B.M., Baumann, M.H., Walther, D., Jimenez-Morigosa, C., Sulima, A., Rice, K.C., Collins, G.T., 2018. The abuse-related effects of pyrrolidine-containing cathinones are related to their potency and selectivity to inhibit the dopamine transporter. Neuropsychopharmacology: official publication of the American College of Neuropsychopharmacology 43(12), 2399–2407.

Gannon, B.M., Galindo, K.I., Mesmin, M.P., Sulima, A., Rice, K.C., Collins, G.T., 2017a. Relative reinforcing effects of second-generation synthetic cathinones: Acquisition of self-administration and fixed ratio dose-response curves in rats. Neuropharmacology.

Gannon, B.M., Rice, K.C., Collins, G.T., 2017b. Reinforcing effects of abused ‘bath salts’ constituents 3,4-methylenedioxypyrovalerone and alpha-pyrrolidinopentiophenone and their enantiomers. Behavioural pharmacology 28(7), 578–581.

Gilpin, N.W., Wright, M.J., Jr., Dickinson, G., Vandewater, S.A., Price, J.U., Taffe, M.A., 2011. Influences of activity wheel access on the body temperature response to MDMA and methamphetamine. Pharmacology, biochemistry, and behavior 99(3), 295–300.

Haughey, H.M., Fleckenstein, A.E., Metzger, R.R., Hanson, G.R., 2000. The effects of methamphetamine on serotonin transporter activity: role of dopamine and hyperthermia. Journal of neurochemistry 75(4), 1608–1617.

Huang, P.-K., Aarde, S.M., Dickerson, T.J., Taffe, M.A., 2012. Acute effects of d-methamphetamine, 3,4-methylenedioxypyrovalerone, 3,4-methylenedioxymethamphetamine, and 4-methylmethcathinone on wheel activity in rats. The FASEB Journal 26(1_MeetingAbstracts), 1040.1043.

Huang, P.K., Aarde, S.M., Angrish, D., Houseknecht, K.L., Dickerson, T.J., Taffe, M.A., 2012. Contrasting effects of d-methamphetamine, 3,4-methylenedioxymethamphetamine, 3,4-methylenedioxypyrovalerone, and 4-methylmethcathinone on wheel activity in rats. Drug and alcohol dependence 126(1-2), 168–175.

Hubner, C.B., Bird, M., Rassnick, S., Kornetsky, C., 1988. The threshold lowering effects of MDMA (ecstasy) on brain-stimulation reward. Psychopharmacology 95(1), 49–51.

Huskinson, S.L., Naylor, J.E., Townsend, E.A., Rowlett, J.K., Blough, B.E., Freeman, K.B., 2017. Self-administration and behavioral economics of second-generation synthetic cathinones in male rats. Psychopharmacology 234(4), 589–598.

Institoris, L., Arok, Z., Seprenyi, K., Varga, T., Sara-Klausz, G., Keller, E., Toth, R.A., Sala, L., Kereszty, E., Rona, K., 2015. Frequency and structure of stimulant designer drug consumption among suspected drug users in Budapest and South-East Hungary in 2012-2013. Forensic science international 248, 181–186.

Jarbe, T.U., Lamb, R.J., Liu, Q., Makriyannis, A., 2003. (R)-Methanandamide and delta9-tetrahydrocannabinol-induced operant rate decreases in rats are not readily antagonized by SR-141716A. Eur J Pharmacol 466(1-2), 121–127.

Javadi-Paydar, M., Harvey, E.L., Grant, Y., Vandewater, S.A., Creehan, K.M., Nguyen, J.D., Dickerson, T.J., Taffe, M.A., 2018a. Binge-like acquisition of alpha-pyrrolidinopentiophenone (alpha-PVP) self-administration in female rats. Psychopharmacology 235(8), 2447–2457.

Javadi-Paydar, M., Nguyen, J.D., Vandewater, S.A., Dickerson, T.J., Taffe, M.A., 2018b. Locomotor and reinforcing effects of pentedrone, pentylone and methylone in rats. Neuropharmacology 134(Pt A), 57–64.

Keel, F., 2019, Tallahassee Police arrest drug trafficker. WCTV. WCTV. https://www.wctv.tv/content/news/Tallahassee-Police-arrest-drug-trafficker-558858941.html

Kenny, P.J., Chen, S.A., Kitamura, O., Markou, A., Koob, G.F., 2006. Conditioned withdrawal drives heroin consumption and decreases reward sensitivity. J Neurosci 26(22), 5894–5900.

Kiyatkin, E.A., Kim, A.H., Wakabayashi, K.T., Baumann, M.H., Shaham, Y., 2015. Effects of Social Interaction and Warm Ambient Temperature on Brain Hyperthermia Induced by the Designer Drugs Methylone and MDPV. Neuropsychopharmacology: official publication of the American College of Neuropsychopharmacology 40(2), 436–445.

Kolanos, R., Sakloth, F., Jain, A.D., Partilla, J.S., Baumann, M.H., Glennon, R.A., 2015. Structural Modification of the Designer Stimulant alpha-Pyrrolidinovalerophenone (alpha-PVP) Influences Potency at Dopamine Transporters. ACS chemical neuroscience 6(10), 1726–1731.

Kornetsky, C., Esposito, R.U., McLean, S., Jacobson, J.O., 1979. Intracranial self-stimulation thresholds: a model for the hedonic effects of drugs of abuse. Archives of general psychiatry 36(3), 289–292.

Lukas, S.E., Mello, N.K., Bree, M.P., Mendelson, J.H., 1988. Differential tolerance development to buprenorphine-, diprenorphine-, and heroin-induced disruption of food-maintained responding in macaque monkeys. Pharmacology, biochemistry, and behavior 30(4), 977–982.

Malberg, J.E., Seiden, L.S., 1998. Small changes in ambient temperature cause large changes in 3,4-methylenedioxymethamphetamine (MDMA)-induced serotonin neurotoxicity and core body temperature in the rat. J Neurosci 18(13), 5086–5094.

Markou, A., Koob, G.F., 1992. Construct validity of a self-stimulation threshold paradigm: effects of reward and performance manipulations. Physiology & behavior 51(1), 111–119.

Matsuta, S., Shima, N., Kakehashi, H., Kamata, H., Nakano, S., Sasaki, K., Kamata, T., Nishioka, H., Miki, A., Zaitsu, K., Tsuchihashi, H., Katagi, M., 2018. Metabolism of alpha-PHP and alpha-PHPP in humans and the effects of alkyl chain lengths on the metabolism of alpha-pyrrolidinophenone-type designer drugs. Forensic Toxicol 36(2), 486–497.

McGee, K., 2019, Bath salts, crack cocaine seized by Columbus County deputies after undercover investigation. WECT 6 News. GRAY TELEVISION LICENSEE, LLC. https://www.wect.com/2019/07/12/bath-salts-crack-cocaine-seized-by-columbus-county-deputies-after-undercover-investigation/

Milian, J., 2015, Riviera Beach Police: Man smoking flakka wielded machete at stepdad. PalmBeachPost.com. Palm Beach Post. http://www.palmbeachpost.com/news/news/crime-law/riviera-beach-police-man-smoking-flakka-wielded-ma/nmW5k/

Miller, M.L., Creehan, K.M., Angrish, D., Barlow, D.J., Houseknecht, K.L., Dickerson, T.J., Taffe, M.A., 2013a. Changes in ambient temperature differentially alter the thermoregulatory, cardiac and locomotor stimulant effects of 4-methylmethcathinone (mephedrone). Drug and alcohol dependence 127(1-3), 248–253.

Miller, M.L., Moreno, A.Y., Aarde, S.M., Creehan, K.M., Vandewater, S.A., Vaillancourt, B.D., Wright, M.J., Jr., Janda, K.D., Taffe, M.A., 2013b. A methamphetamine vaccine attenuates methamphetamine-induced disruptions in thermoregulation and activity in rats. Biological psychiatry 73(8), 721–728.

Nagy, E.K., Overby, P.F., Olive, M.F., 2020. Reinforcing Effects of the Synthetic Cathinone α-Pyrrolidinopropiophenone (α-PPP) in a Repeated Extended Access Binge Paradigm. Frontiers in Psychiatry 11(862).

National Research Council (U.S.). Committee for the Update of the Guide for the Care and Use of Laboratory Animals., Institute for Laboratory Animal Research (U.S.), National Academies Press (U.S.), 2011. Guide for the care and use of laboratory animals: Eigth Edition, 8th ed. National Academies Press,, Washington, D.C., pp. xxv, 220 p.

Nelson, K.H., Hempel, B.J., Clasen, M.M., Rice, K.C., Riley, A.L., 2017. Conditioned taste avoidance, conditioned place preference and hyperthermia induced by the second generation ‘bath salt’ alpha-pyrrolidinopentiophenone (alpha-PVP). Pharmacology, biochemistry, and behavior 156, 48–55.

Nelson, K.H., Manke, H.N., Imanalieva, A., Rice, K.C., Riley, A.L., 2019. Sex differences in alpha-pyrrolidinopentiophenone (alpha-PVP)-induced taste avoidance, place preference, hyperthermia and locomotor activity in rats. Pharmacology, biochemistry, and behavior 185, 172762.

Nguyen, J.D., Aarde, S.M., Cole, M., Vandewater, S.A., Grant, Y., Taffe, M.A., 2016. Locomotor Stimulant and Rewarding Effects of Inhaling Methamphetamine, MDPV, and Mephedrone via Electronic Cigarette-Type Technology. Neuropsychopharmacology: official publication of the American College of Neuropsychopharmacology 41(11), 2759–2771.

Nguyen, J.D., Bremer, P.T., Hwang, C.S., Vandewater, S.A., Collins, K.C., Creehan, K.M., Janda, K.D., Taffe, M.A., 2017. Effective active vaccination against methamphetamine in female rats. Drug and alcohol dependence 175, 179–186.

Nguyen, J.D., Grant, Y., Taffe, M.A., 2019. Paradoxical changes in brain reward status during opioid self-administration in a novel test of the negative reinforcement hypothesis. bioRxiv, 460048.

Richman, E., Skoller, N.J., Fokum, B., Burke, B.A., Hickerson, C.A., Cotes, R.O., 2018. alpha-Pyrrolidinopentiophenone (“Flakka”) Catalyzing Catatonia: A Case Report and Literature Review. Journal of addiction medicine.

Schindler, C.W., Thorndike, E.B., Walters, H.M., Walther, D., Rice, K.C., Baumann, M.H., 2019. Stereoselective neurochemical, behavioral, and cardiovascular effects of alpha-pyrrolidinovalerophenone enantiomers in male rats. Addiction biology, e12842.

Segal, D.S., Kuczenski, R., 1999. Escalating dose-binge stimulant exposure: relationship between emergent behavioral profile and differential caudate-putamen and nucleus accumbens dopamine responses. Psychopharmacology 142(2), 182–192.

Simmler, L., Buser, T., Donzelli, M., Schramm, Y., Dieu, L.H., Huwyler, J., Chaboz, S., Hoener, M., Liechti, M., 2013. Pharmacological characterization of designer cathinones in vitro. British journal of pharmacology 168(2), 458–470.

Simmler, L.D., Rickli, A., Hoener, M.C., Liechti, M.E., 2014. Monoamine transporter and receptor interaction profiles of a new series of designer cathinones. Neuropharmacology 79, 152–160.

Swirko, C., 2020, Woman who claimed self-defense in shooting gets drug charge. The Gainesville Sun. Gannett Media Corp. https://www.gainesville.com/news/20200424/woman-who-claimed-self-defense-in-shooting-gets-drug-charge

Taffe, M.A., Creehan, K.M., Vandewater, S.A., 2015. Cannabidiol fails to reverse hypothermia or locomotor suppression induced by Delta(9) - tetrahydrocannabinol in Sprague-Dawley rats. British journal of pharmacology 172(7), 1783–1791.

WALBNewsTeam, 2020, Valdosta police make drug arrest. WALB 10 News. GRAY TELEVISION LICENSEE, LLC. https://www.walb.com/2020/05/11/valdosta-police-make-drug-arrest/

Watterson, L.R., Kufahl, P.R., Nemirovsky, N.E., Sewalia, K., Grabenauer, M., Thomas, B.F., Marusich, J.A., Wegner, S., Olive, M.F., 2014. Potent rewarding and reinforcing effects of the synthetic cathinone 3,4-methylenedioxypyrovalerone (MDPV). Addiction biology 19(2), 165–174.

Wise, J., 2016, The Obscure, Legal Drug That Fuels John McAfee. New York Magazine Daily Intelligencer. New York Media. http://nymag.com/daily/intelligencer/2016/09/the-obscure-legal-drug-that-fuels-john-mcafee.html

Wright, M.J., Jr., Angrish, D., Aarde, S.M., Barlow, D.J., Buczynski, M.W., Creehan, K.M., Vandewater, S.A., Parsons, L.H., Houseknecht, K.L., Dickerson, T.J., Taffe, M.A., 2012. Effect of ambient temperature on the thermoregulatory and locomotor stimulant effects of 4-methylmethcathinone in Wistar and Sprague-Dawley rats. PloS one 7(8), e44652.

